# Comparison Of Multi-locus Sequence Typing software For next generation sequencing data

**DOI:** 10.1101/117770

**Authors:** Andrew J. Page, Nabil-Fareed Alikhan, Heather A. Carleton, Torsten Seemann, Jacqueline A. Keane, Lee S. Katz

## Abstract

Multi-locus sequence typing (MLST) is a widely used method for categorising bacteria. Increasingly MLST is being performed using next generation sequencing data by reference labs and for clinical diagnostics. Many software applications have been developed to calculate sequence types from NGS data; however, there has been no comprehensive review to date on these methods. We have compared six of these applications against real and simulated data and present results on: 1. the accuracy of each method against traditional typing methods, 2. the performance on real outbreak datasets, 3. in the impact of contamination and varying depth of coverage, and 4. the computational resource requirements.

**DATA SUMMARY:**

1. Simulated reads for datasets testing coverage and mixed samples have been deposited in Figshare; DOI: https://doi.org/10.6084/m9.figshare.4602301.vl
2. Outbreak databases are available from Github; url - https://github.com/WGS-standards-and-analysis/datasets
3. Docker containers used to run each of the applications are available from Github; url – https://tinyurl.com/z7ks2ft
4. Accession numbers for the data used in this paper are available in the Supplementary material.

**We confirm all supporting data, code and protocols have been provided within the article or through supplementary data files. ☒**

**IMPACT STATEMENT:** Sequence typing is rapidly transitioning from traditional sequencing methods to using whole genome sequencing. A number of *in silico* prediction methods have been developed on an *ad hoc* basis and aim to replicate Multi-locus sequence typing (MLST). This is the first study to comprehensively evaluate multiple MLST software applications on real validated datasets and on common simulated difficult cases. It will give researchers a clearer understanding of the accuracy, limitations and computational performance of the methods they use, and will assist future researchers to choose the most appropriate method for their experimental goals.

## INTRODUCTION

A small number of bacterial foodborne pathogens, such as *Salmonella, Campylobacter, Listeria* and *Escherichia*, cause a huge burden of disease in humans and animals. The most deadly serovars of *Salmonella* cause over 230,000 deaths every year (1) with increasing instances of multiple drug resistance emerging (2). With *L. monocytogenes*, although the case count is small, the case-fatality rate is high at approximately 21% to 38% (3,4). Additionally, the economic burden is high (5). In the US, each foodborne illness can cost anywhere from hundreds to millions of US dollars depending on the organism. Therefore, investigating potential foodborne outbreaks and preventing any illness is advantageous from both economic and public health standpoints. In order to understand these bacteria in more depth, there have been many studies to describe their population structure using phylogenetic methods based on multi-locus sequence typing (MLST) (6,7).

Additionally, there are many large-scale efforts to surveil these pathogens. One of the most successful programs has been PulseNet International (8) which aids in the detection of common source outbreaks. Historically, PulseNet International has used pulsed-field gel electrophoresis (PFGE) to find genomic matches among bacterial isolates. Recently, large numbers of isolates are being whole genome sequencing (WGS) through an initiative between The Centers for Disease Control and Prevention (CDC), The US Food and Drug Administration (FDA), The US Department of Agriculture (USDA), and The National Center for Biotechnology Information (NCBI). Through this collaboration, every *L. monocytogenes* genome that is discovered in the food supply, or in clinical samples, is being sequenced and uploaded to NCBI's Sequence Read Archive (SRA) database. This collaboration has since started sequencing a large percentage of *E. coli, S. enterica, C. coli, C. jejuni*, and many others with the eventual goal of completely switching from PFGE to WGS. In Europe, Public Health England (PHE) sequences every *Salmonella* and *Mycobacterium tuberculosis* isolate submitted to them and deposit the data in SRA. Perceiving a future need for worldwide collaboration on these new methods, the Global Microbial Identifier (GMI) (9) partnership was initiated in 2013 to encourage data sharing among all nations for many purposes including public health and research. As of February 2017, at least 43 nations have participated in GMI, indicating a growing interest in WGS methodology and data sharing. GMI continues to enhance the value of the sequence data by encouraging submitters to provide rich, useful metadata to accompany the deposited samples. By switching from classic methods, such as Sanger sequencing or PFGE, reference labs gain richer, more valuable insights for minimal additional cost.

To aid in population structure studies and in epidemiological investigations, multi-locus sequence typing (MLST) has been used for nearly two decades (10) to categorise different clonal expansions of these pathogens into broad categories, based on allelic variation amongst seven highly conserved housekeeping genes. Sequence typing can be performed using both next generation sequencing (NGS) and classical sequencing techniques. A number of software applications have been developed using a variety of fundamentally different techniques to calculate sequence types from NGS data. However, there has been no comprehensive review to date on the accuracy, computational performance, robustness and ease of use of these methods. In this paper we evaluate multiple MLST software applications on a variety of datasets, both real and simulated such as: 1.) standard sets of outbreak data from the Gen-FS WGS Standards and Analysis working group (11,12) which includes *Campylobacter jejuni, Escherichia coli, Listeria monocytogenes* and *Salmonella enterica*, 2.) *Salmonella* isolates which have been typed using both traditional capillary sequencing and next generation sequencing, 3.) simulated reads of varying coverage, and 4.) simulated mixed strains. Here, we describe a comprehensive list of command-line tools for MLST analysis and benchmark them with these standardised datasets in terms of accuracy and computer resources required.

## SOFTWARE OVERVIEW

MLST software can be categorised according to the input data they accept; there are tools that use raw sequence reads and tools that use *de novo* assemblies. Calling MLST from raw reads avoids the need to fully reconstruct the whole genome, theoretically allowing for a lower running time. However in practice *de novo* assembly is routinely performed for bacteria (13) and assemblies may already be available for any given MLST analysis leading to faster sequence typing. The process of *de novo* assembly can introduce artefacts, particularly from short reads. For example, a gene may be fragmented over multiple contigs. A full overview is given in Table 1. In general, the desired characteristics of MLST software include:

1. high specificity of calling sequence types (STs),
2. resilient in the face of mixed samples,
3. tolerant with low sequencing coverage,
4. efficient in computational and disk resources,
5. packaged for easy dependency management and installation,
6. validated with automated tests to verify functionality works as intended,
7. open source,
8. and scalability to large numbers of isolates.

**Table 1:** Overview of MLST software.

| Software | Input | Algorithm | Licence | Availability | Tests | Installation | Interface |
| --- | --- | --- | --- | --- | --- | --- | --- |
| Ariba (23) | reads | Assembly | GPL3 | Github | Yes | pip, apt, docker | Command line |
| BigsDB (15) | contigs | BLASTn | GPL3 | Github | No | Manual | Website |
| BioNumerics (21) | NA | NA | Bespoke | Proprietary | NA | NA | NA |
| CGE-MLST (28) | contigs | BLASTn | Apache 2 | Bitbucket | No | docker | Command line / Website |
| Enterobase (14) | reads | uBLAST/ BLASTn | NA | NA | NA | NA | Website |
| lyve-MLST (27) | contigs | BLAST | MIT | Github | No | Manual | Command line |
| MOST (18) | reads | Mapping | FreeBSD | Github | No | Manual | Command line |
| MLSTcheck (25) | contigs | BLASTn | GPL3 | Github | Yes | CPAN, docker | Command line |
| mlst (24) | contigs | BLASTn | GPL2 | Github | No | Brew | Command line |
| stringMLST (20) | reads | *k*-mer | Bespoke | Github | No | Manual | Command line |
| SRST2 (16) | reads | Mapping | BSD | Github | Yes | apt, pip | Command line |

The interfaces to the software applications fall into two categories, those that operate on the command line and those that have a graphical interface. Command line input allows for high throughput analysis, but have a high barrier to entry for non-technical users. Graphical interfaces, such as websites, provide point and click interfaces which non-technical users find easier to use initially; however they are often limited to the analysis of a few samples at a time. To reduce the impact of this limitation, some websites precompute results by downloading raw data directly from the short read archives (14,15).

Most of the software packages are available under open source licences, with source code available in public repositories such as GitHub (https://github.com). Source code availability facilitates transparency for the underlying methods. Comprehensive automated tests, if designed correctly, ensure stability within software applications. Applications packaged for easy installation and dependency management such as: Apt (debian), Homebrew, Docker, PyPy, and CPAN allow for the software to be installed in one step, allowing for immediate use by a range of users. An overview of MLST software applications follows.

*Short Read Sequence Typing 2 (SRST2)* (16) takes raw reads as input on the command line and uses a mapping based approach to align reads to the alleles. It is packaged for easy dependency installation and is widely used for a variety of applications in addition to MLST including: antibiotic resistance prediction, virulence gene detection and serotyping (17). The software licence is one of the most liberal of all and it has unit tests.

*Metric-Oriented Sequence Typer (MOST)* (18) builds upon *SRST* (version 1)(19) and uses a mapping based approach to align alleles to reads, with a traffic light system indicating the confidence in the ST calling. One major difference to SRST2 is that it takes a 100 base flanking region around the locus from a reference genome, reducing the impact of coverage drop off at the ends of the sequences. Additionally it can assign predicted serovars to Salmonella isolates. It is used by Public Health England on clinical isolates and has strict, well defined conservative criteria for calling sequence types to ensure accuracy.

*stringMLST* (20) takes raw reads as input on the command line and uses *k*-mers to detect MLST alleles. Instead of detecting allele coverage or parsing for potential SNPs, an allele call is made by identifying the allele with the most number of matching *k*-mers. The use of *k*-mers gives a substantial speed advantage, but at the expense of accuracy. This method is fast enough to detect sequence types in real time during sequencing, so holds much promise for the future. It is free for non-commercial purposes and it has no automated tests.

*BioNumerics* (21) from Applied Maths is a commercial application which is widely used by public health labs to calculate sequence types. Due to its proprietary nature a full review is not possible, however the authors describe a *k*-mer based sequence typing method in a patent (22).

*Antibiotic Resistance Identification By Assembly* (*ARIBA*) (23) takes raw reads as input on the command line and uses a combination of mapping and local *de novo* assembly to calculate alleles. Like SRST2 it can be used more generally for gene detection and classification, allowing for antibiotic resistance prediction, virulence gene detection and plasmid replication gene classification. It is open source, has extensive unit tests and is packaged for easy installation.

*mlst* (24) takes *de novo* assemblies as input on the command line and uses BLASTn to align sequences to alleles. It is very fast and searches all databases on pubMLST to automatically detect the organism, then calculates the ST. Installation is very easy using *brew*.

*MLSTcheck* (25) takes *de novo* assemblies as input on the command line and uses BLASTn to align sequences to alleles. It is packaged for easy dependency installation, and has unit test coverage. It produces a multi-FASTA alignment of concatenated allele sequences for each sample, which allows for phylogenetic trees to be easily constructed.

*EnteroBase* (14) is a website which incorporates sequencing data from both public databases and directly from users for four genera *(Salmonella, Escherichia, Yersinia* & *Moraxella)*, and assembles it *de novo* with an adjusted pipeline using SPAdes (26). It is maintained by the Achtman group, who have been involved with MLST since the beginning (10). Alleles are called using uBLAST and BLASTp, which allows for high sensitivity for divergent allele variants. However, the source code is not publicly available.

Bacterial Isolate Genome Sequence Database (BigsDB) (15) is a web service whose primarily purpose is the management of sequence typing databases, as opposed to querying them. It is used by the majority of schemes as the backend for storing their typing data. The database can be queried in two ways, via a web interface, or programmatically through a REST API. There is no described command line interface for queries, however the mechanisms are in place to allow for it in the future.

lyve-MLST (27) is a wrapper around the Institut Pasteur web service for calculating MLST. It is hard coded to work with *Listeria monocytogenes*, does not have versioning and is no longer under active development.

*MLST* from the Center for Genomic Epidemiology (MLST-CGE) (28) is a web based method for calculating MLST. It can take assembled genomes or raw sequencing reads. If raw sequencing reads are provided, it performs a de novo assembly. Alleles are called using a BLAST based method.

## DATABASE AVAILABILITY

The availability of databases containing alleles and ST profiles for different species is an important aspect of any MLST software application as outlined in Table 2, since this dictates how easy it is to use the software. These databases also need to be kept up to date as the underlying schemes are constantly being extended as new isolates are sequenced. Out of date databases can mean that rapidly emerging clonal expansions may be missed, impairing epidemiological investigations. *ARIBA, mlst, MLSTcheck, stringMLST* and *SRST2* all provide automated scripts to download all of the latest databases from pubMLST (15), which are immediately ready to use. This provides immediate access to schemes for over 125 species. *mlst* and *stringMLST* go one step further and additionally bundles all available databases in their software repository, which are regularly updated. *MOST* do not provide an automated method for downloading new or updated databases, instead directing researchers to a set of manual steps. They do provide a small number of bundled databases (6 and 9 respectively) however these only represent a fraction of the currently available databases on pubMLST. The databases bundled with *MOST* were last updated in December 2015, so are missing all recent updates and additions to the schemes, including new STs, so researchers cannot be certain novel results are indeed novel.

**Table 2:** Overview of the MLST databases available with each software application.

| Software | Automated downloads | Bundled databases | Age of bundled DBs<sup>1</sup> | DBs ready to use |
| --- | --- | --- | --- | --- |
| Ariba | Yes | 0 | - | Yes |
| mlst | Yes | 125 | 1 month | Yes |
| MLSTcheck | Yes | 0 | - | Yes |
| MOST | No | 6 | >1 year | Yes |
| SRST2 | Yes | 0 | - | Yes |
| stringMLST | Yes | 128 | 1 month | Yes |

<sup>1</sup>The age of the bundled databases was calculated on the 15 March 2017.

## EVALUATION

A meaningful comparison could only be performed with the six open source command line MLST software applications, *ARIBA* (v2.7.2), *mlst* (v2.8), *MLSTcheck* (v2.1.1630910), MOST (v 2e3da07), *SRST2* (v0.2.0) and *stringMLST* (v0.3.6). *Lyve-MLST* was excluded because it was hard coded to work with a single database. *BigsDB* and *Enterobase* were excluded as they are web services with extensively featured pipelines and the computational performance of the MLST calling component could not be measured independently. *MLST-CGE* was excluded because an essential internally hosted software repository was unavailable at the time of testing. Partial results are available for *Enterobase* for some datasets, where relevant.

Each application was evaluated on 4 different datasets, 2 real and 2 simulated. Dataset 1 contains 85 samples from standard sets of outbreak data from the Gen-FS WGS Standards and Analysis working group (11). Dataset 2 consists of 72 *Salmonella* samples which have been sequenced using traditional capillary sequencing and using Illumina NGS technologies. This allows for technology independent validation. Dataset 3 consists of artificially generated reads with varying levels of coverage. From this the minimum sequence depth required for each software application can be calculated. Dataset 4 consists of artificially generated reads from 2 different *Salmonella* serovars where all alleles differ, mixed in different ratios out of a total depth of coverage of 50X. The accuracy of applications can then be determined with mixed samples (a common case) and the point at which the mixed samples become detectable.

The experiments for Dataset 1 were performed using the CDC compute infrastructure. For the rest of the experiments (2, 3, 4) we used the MRC CLIMB OpenStack cloud (29) as the base platform for the evaluations. Each of the applications was run in their own Docker container (30) available from (31). The Debian Testing distribution was used as the base operating system for all containers as it provides access to a large range of up-to-date bioinformatics software. The host VM had 4 cores and 32GB of RAM running Ubuntu 16.04 (LTS), however only a single core was used for the evaluations. All datasets used for this analysis are available for download as described in the data bibliography or from the public archives using the accession numbers in the Supplementary Material. Where assemblies were required as input to MLST applications, the raw reads were *de novo* assembled with SPAdes (v3.9.0) (26) using the default parameters. SPAdes was chosen as it is widely used and consistently produces high quality results on bacterial data (32).

## REAL OUTBREAK DATASET

Standard datasets (11) covering *Listeria monocytogenes* from stone fruit (33), *Escherichia coli* from sprouts (34), *Campylobacter jejuni* from raw milk (35) and *Salmonella enterica* from spicy tuna (36) comprising of 85 samples were analysed by each of the software applications. These are real outbreak datasets where there was substantive epidemiological investigations and full details can be found in (12). No false positives were reported by any application, they either made the correct call, a low confidence call or no call. A summary of the overall performance is provided in Table 3, with extended details available in Table S1. There was a wide variation in the results, with only 2 applications (*stringMLST*, and *MLSTcheck)* correctly calling all of the STs. *MOST* failed to confidently call any of the spicy tuna *Salmonella* samples, but did identify the correct STs, flagged as low confidence (amber). There was a 29-fold variation in the running times between the applications (*stringMLST* vs *SRST2*) using raw reads as input (Table 1). This extra computation imposes financial costs and increases the analysis time after sequencing.

**Table 3:** Summary of performance of each algorithm on real outbreak data for 4 different species (85 samples).

| Software | Total time (m) | Correct ST (%) | No call/low confidence (%) |
| --- | --- | --- | --- |
| ARIBA | 109.5 | 98.8 | **1.2** |
| mlst<sup>1</sup> | 1.9 (+ 2873) | 96.5 | 3.5 |
| MOST<sup>2</sup> | 1189.7 | 49.4 | 50.6 |
| MLSTcheck<sup>1</sup> | 63.8 (+ 2873) | **100** | 0 |
| SRST2 | 2380.2 | 95.3 | 4.7 |
| stringMLST | **80.8** | **100** | 0 |

<sup>1</sup>The time to assemble with SPAdes before running the applications was 2873 minutes and is included separately. <sup>2</sup>MOST identified the correct ST in 97.6% of cases, but flags 48.2% of these calls as low confidence.

## COMPARISON TO CAPILLARY DATA

This dataset consists of 72 *Salmonella* samples which have been sequenced using traditional capillary sequencing (originally deposited in http://mlst.warwick.ac.uk) and sequenced using NGS. This allows for technology independent validation of the NGS MLST software applications. The samples cover a wide range of *Salmonella*, with hosts including humans, reptiles, birds, farm/domestic animals, and environmental, collected between 1940-2014. The dataset contains an estimated 32 different STs, with 38 of the samples predicted to have a serovar of Typhimurium, which causes severe disease in a wide range of hosts, including human. Full details of the samples (including accessions) and results are in Table S2. The ST calls match in 89% (64/72) of cases between the capillary data and the NGS MLST software applications, which additionally includes MLST results from the Enterobase website. Two samples (RKS1252 and RKS1256) are suspected sample swaps with each other. The sample E698 differs between the trace results and all other methods with no overlapping alleles. It is likely possibly a sample swap with another unknown sample or the original sample contained multiple strains. For OLC-1602 and 556-59/192 6 out of 7 alleles match in all of the results, but the trace data reports a single different allele. Whilst trace data is recognised by the community as a gold standard, it is not error free (37), with calls sometimes made using a single read, leaving little resilience to sequencing errors. As the NGS data has very high depth of coverage (over 30X) of this allele, it is likely that the NGS results are correct. Nearly all of the calls from *MOST* are low confidence (rated amber) however they correlate with the results from the other applications. Four samples are flagged by multiple applications as problematic, however in every case the trace data and *stringMLST* have confidently called an ST, indicating an obvious contaminant has been missed by *stringMLST. MLSTcheck* calls an incorrect ST for 2 samples, getting one of the alleles wrong. Overall we conclude that whilst the MLST results between trace data and NGS data are nearly identical, the MLST based on NGS data is more accurate and reliable when presented with edge cases.

## IMPACT OF DEPTH OF COVERAGE

The impact of depth of coverage over the MLST genes was assessed by artificially generating perfect paired ended reads with a length of 125 bases and a median insert size of 400 with varying levels of coverage using FASTAQ (v3.14.0). The allele sequences plus 500 base flanking regions are extracted from *S*. Typhi CT18 (38), accession AL513382, and artificial paired end reads were generated with average depths of coverage from 1 to 30. The simulated reads were free from sequencing errors to allow for the effect of coverage alone to be measured. Therefore, the minimum effective depth of coverage for each application can be tested. All applications could accurately call STs when the coverage was greater than 12X however below this the minimum depth of coverage applications required varied greatly, as shown in Figure 1. *stringMLST* correctly called the ST with just 3X coverage, however it gave false positive results for lower coverage alleles. *ARIBA* correctly calls the ST from 5X with no false positive results. *SRST2* correctly calls the ST from 12X coverage with no false positive results, however it does correctly identify the ST from 6X with low confidence.

**Figure 1:**
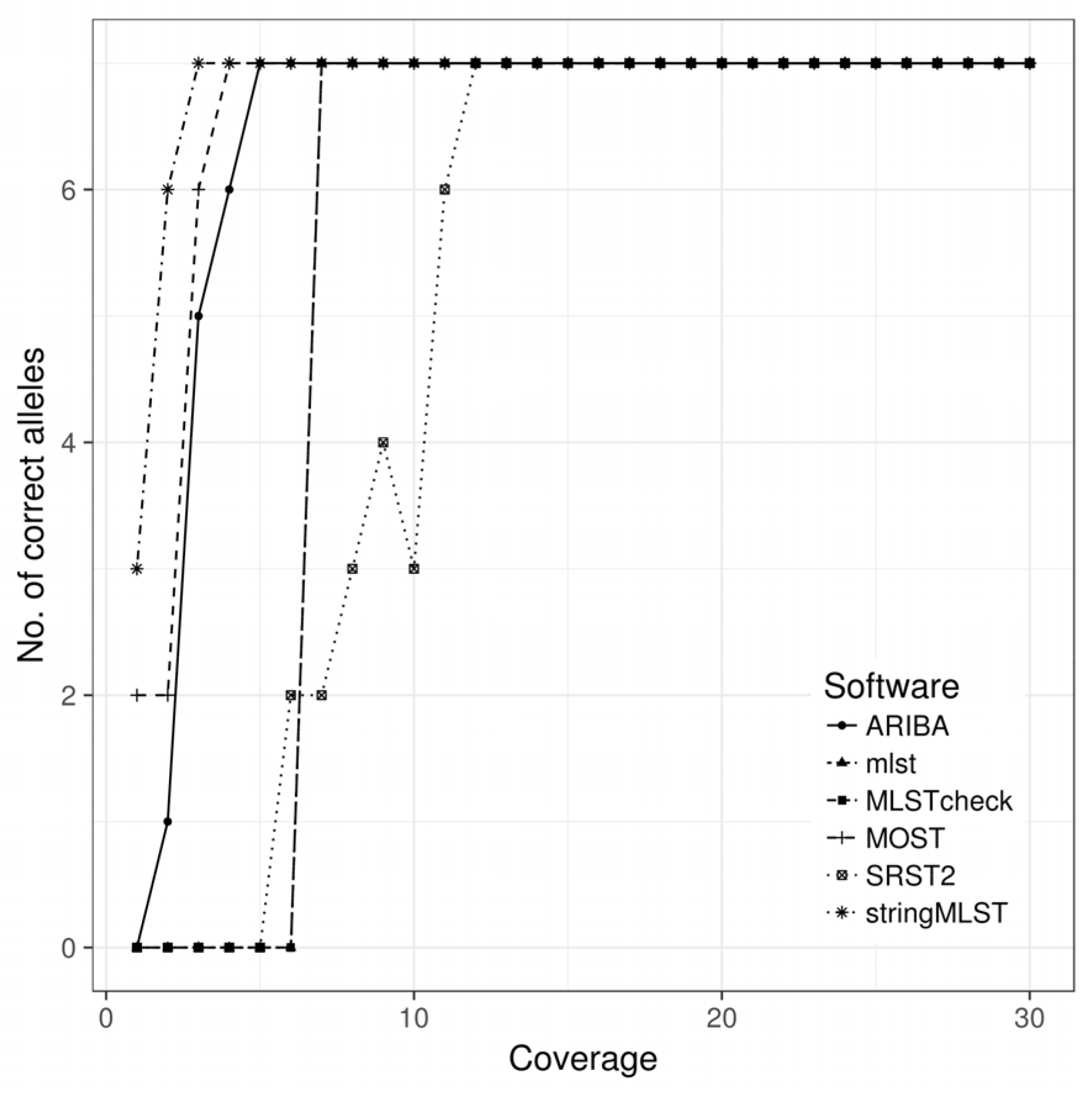
Number of correct calls of each application as coverage increases. Each ST consists of 7 alleles, and all 7 must be correctly & confidently called to calculate a ST.

The computational resources required varied greatly with *stringMLST* taking just 10 seconds to call an ST with 30X coverage, as shown in Figures 2, and the final disk space requirements were negligible as shown in Figure 3. Whilst minimising the disk space resources need for the application is generally positive, *stringMLST* does not output enough information about the allele calls to allow for further analysis, for example, to interrogate a false positive result. The time to call an ST at 30X with *ARIBA* was 40 seconds with 0.1 Mbytes of output data. The disk space requirement is higher than *stringMLST* but provides the allele assemblies used to call the ST which is useful for further analysis. *SRST2* is an order of magnitude slower taking over 500 seconds to call an ST at 30X. The disk space required for the final output is also very substantial at 147 Mbytes which equates to a storage cost of 475 bytes per base of sequencing as shown in Figure 3. While *MOST* confidently correctly calls each individual allele from 4X, the overall ST call is flagged as low confidence below 10X due to its inherently conservative nature. The running time given for *mlst* and *MLSTcheck* includes the de novo assembly time with SPAdes which accounts for most of the running time. *MLSTcheck* takes on average 4 times longer (25 seconds per sample) to return a result than *mlst* (5.9 seconds per sample) with the final results between the two being identical.

**Figure 2:**
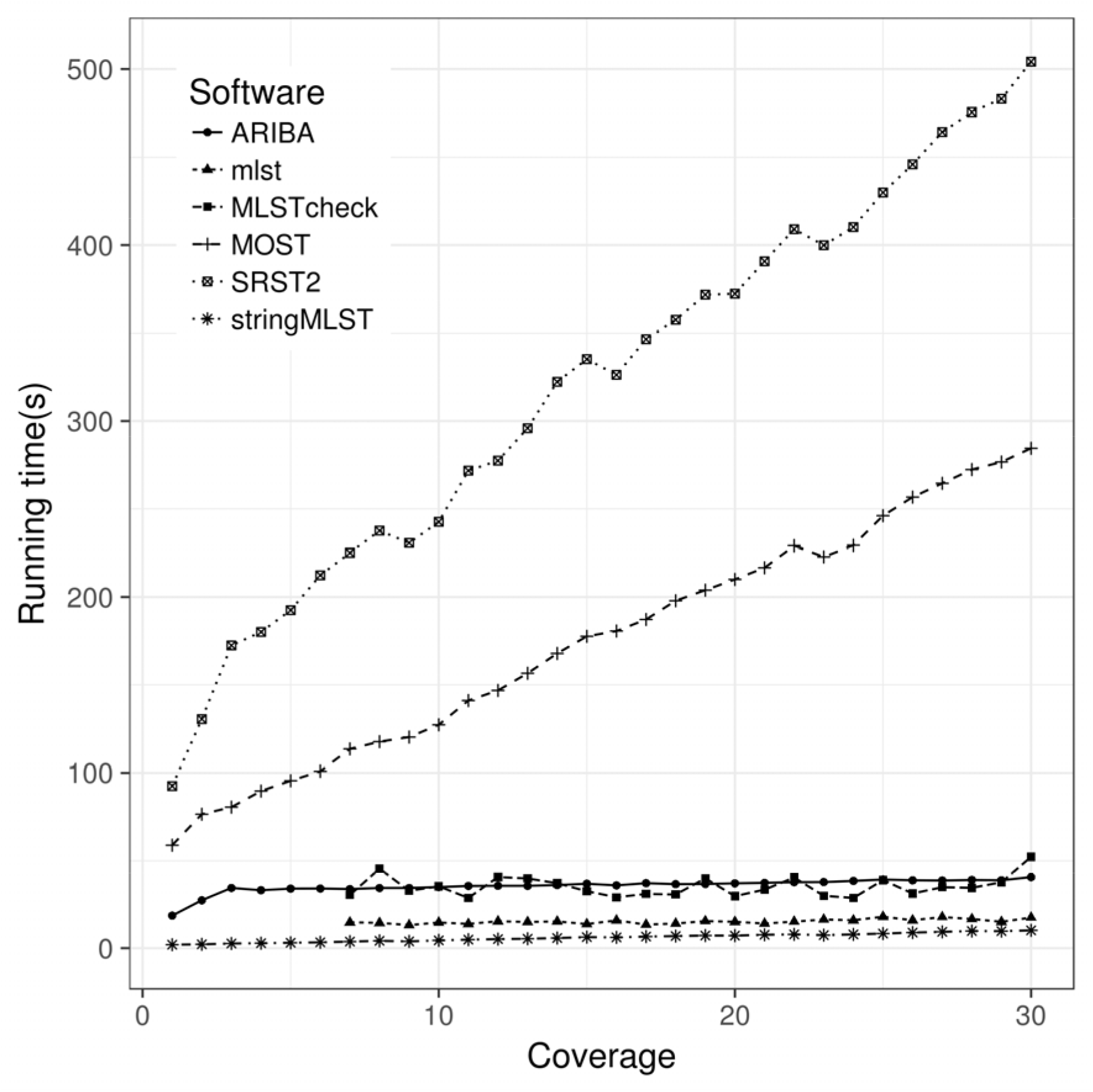
Running time (seconds) of each application as the coverage increases to assess the impact of the depth of coverage. No assembled contiguous sequences could be generated where the coverage was less than 7X, as such no data was recorded for the reliant methods *(mlst & MLSTcheck)*.

**Figure 3:**
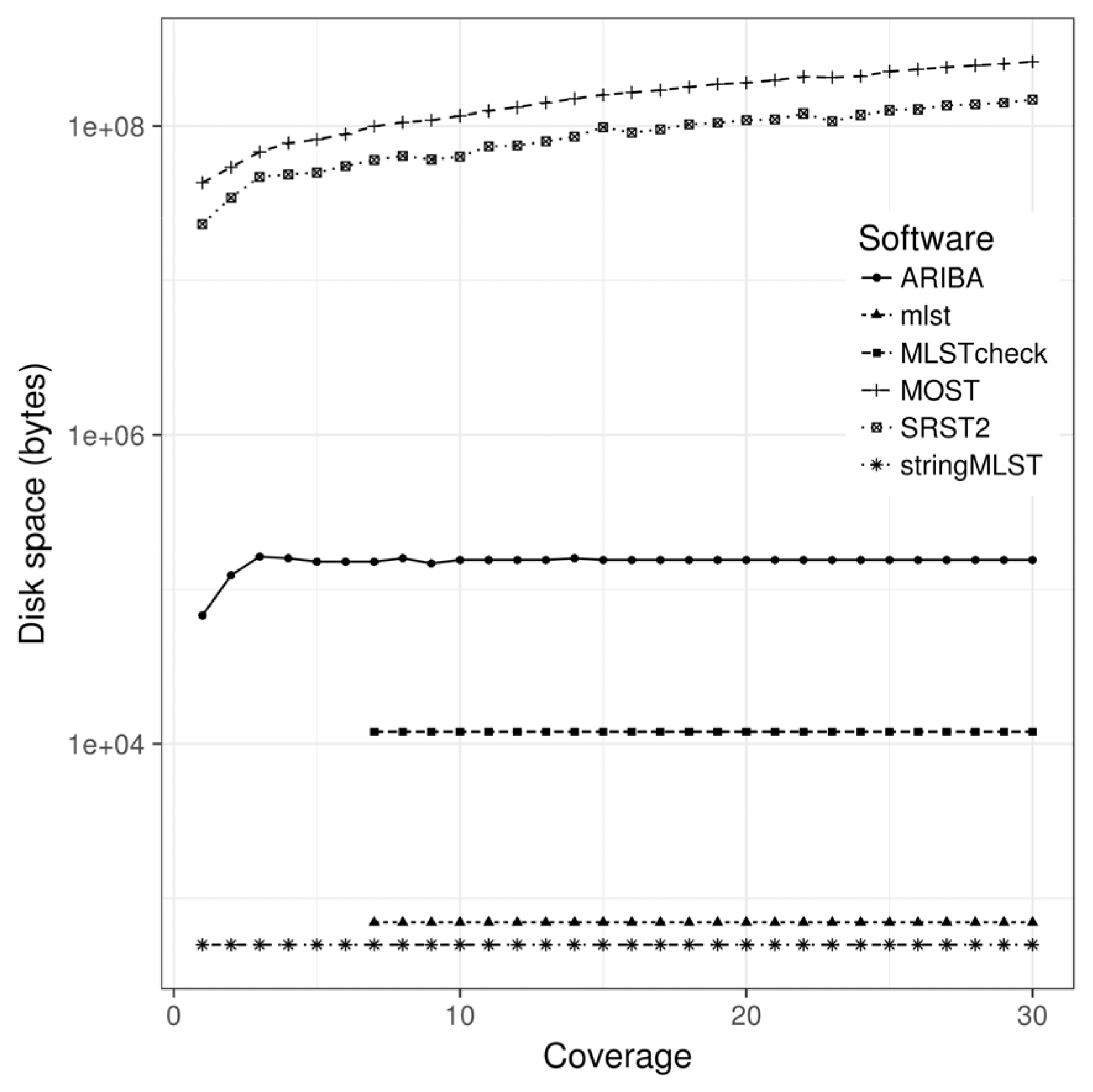
Disk space requirements in bytes for each software application as the depth of coverage increases. Due to the large difference between applications, a log scale is used.

## IMPACT OF MIXED SAMPLES

Contamination and mixed colonies are a standard complexity in microbiology (39). To understand the behaviour of the different MLST software applications in the presence of more than one strain, we constructed a simulated dataset consisting of two *Salmonella* samples with different alleles in varying ratios. This allowed us to see at what point contamination/mixed strains becomes detectable. Once detected we would expect an MLST application to flag the results as low confidence or provide no result at all to avoid false positives. The flip side of this is that if algorithms are too sensitive to low level contamination and sequencing errors, they become less useful on real world applications, so need to be tolerant to some low level noise.

The allele sequences plus 500 base flanking regions were extracted from *S*. Typhi CT18 (38), accession AL513382, and *S*. Weltevreden 10259 (40), accession LN890518. Artificial paired end reads were generated using FASTAQ to give a total coverage of 50X, beginning with CT18 at 1X and 10259 at 49X in a single FASTQ file. The coverage of each sample was varied in steps of 1X to generate a dataset of 49 FASTQ files. Figure 4 shows that the accuracy of the software varies, but follows a general pattern, calling the sample with the highest coverage at the highest levels, with uncertainty in the middle as the proportion of the two samples becomes similar. The worst case is where a software application calls an ST with high confidence which is not in the underlying data (false positive), and only occurs with *stringMLST. MOST* and *ARIBA* is highly conservative detecting that there are mixed samples when the samples are at very low levels of coverage (at 4-5X). *MLSTcheck* and *mlst* both perform identically, with the performance linked to how well SPAdes assembled the underlying genomes. There is no clear boundary with *SRST2* and it varies between high quality calls and low confidence calls as the mixing of the samples changes.

**Figure 4:**
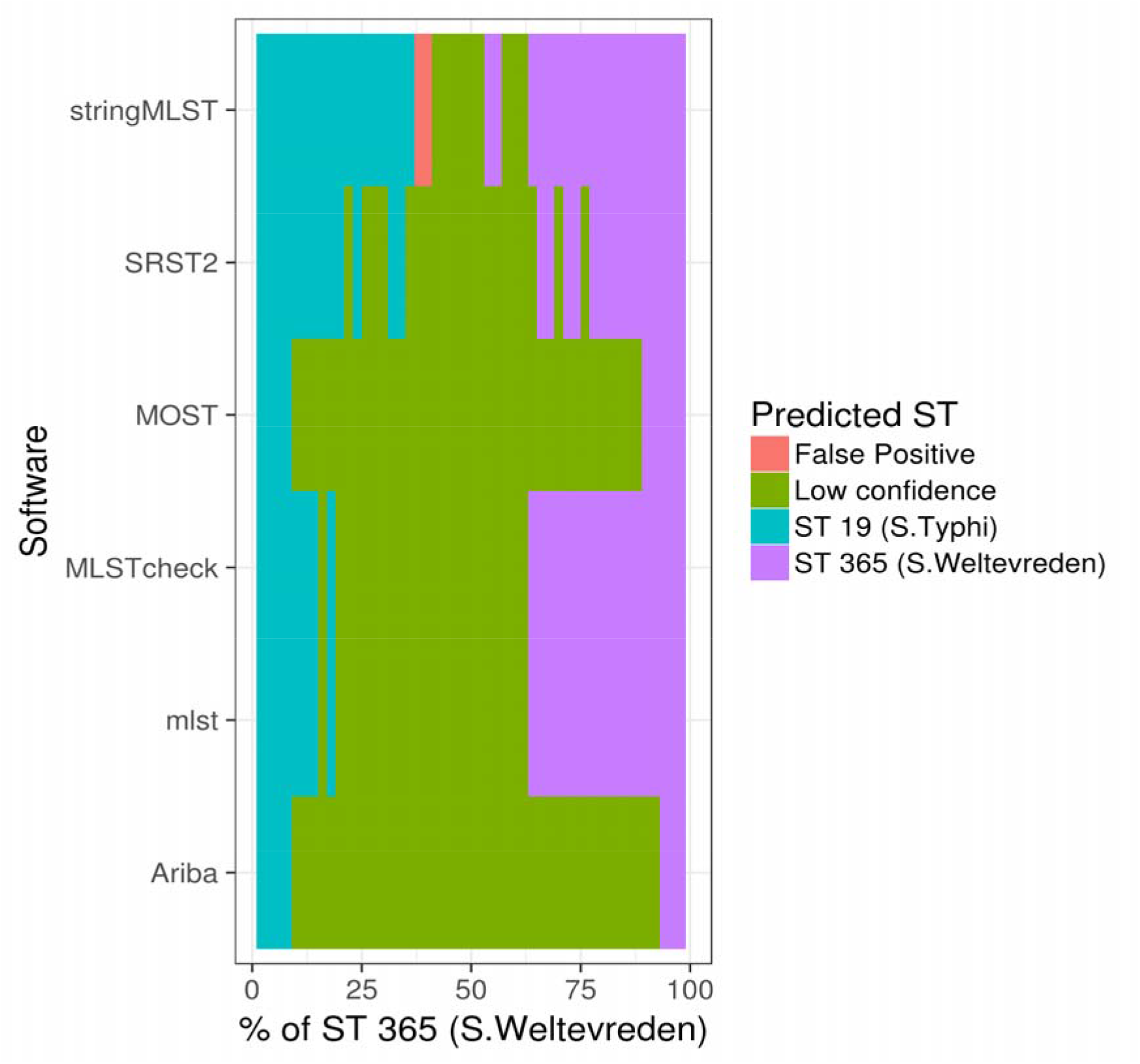
Sequence types called by each software application when given data containing 2 different *Salmonella* samples in varying ratios of abundance. Where there is no ST called, or where the ST has any ambiguity at all, it is marked as ‘low confidence’. A false positive is where an ST is called with high confidence and is not one of the 2 samples in the raw data.

## CONCLUSION

It is clear that applications which have been developed to address this problem have strengths and weaknesses in different areas. Problems with some software include: out of date databases, computationally inefficient methods, false positive results, inability to call alleles at low coverage and variable performance in the presence of mixed samples. Overall though, the accuracy is as good as, or better than, traditional MLST calling methods based on trace sequencing data. The key is to choose the best application for the problem being addressed. When starting from raw reads, for situations where a quick answer is needed and accuracy is not critical, *stringMLST* performs best. Where accuracy and speed are important, *ARIBA* performs the best. Where well defined criteria for calls are required, such as for formal certification of protocols, *MOST* is best suited. Finally, where a *de novo* assembly already exists, *mlst* provides the fastest results.

## AUTHOR STATEMENTS

This work was supported by the Wellcome Trust (grant WT 098051). This work was made possible through support from the Advanced Molecular Detection (AMD) Initiative at the Centers for Disease Control and Prevention. Thanks to João Carriço, Miguel Machado, Anthony Underwood, Martin Flunt, King Jordan and Simon Harris for their helpful discussions and suggestions.

## ABBREVIATIONS

MLST: Multi-locus sequence typing
NGS: Next generation sequencing
PFGE: Pulsed-field gel electrophoresis

